# Directed evolution of laccase from *Thermus thermophilus* HB27 reveals a variant with enhanced high-temperature catalytic activity and electrochemical properties

**DOI:** 10.64898/2026.10.09.757997

**Authors:** Beatriz Miranda-Zaragoza, Paul Gaytán-Colín, Guillermo Huerta-Miranda, Leticia Ramírez-Silva, Margarita Miranda-Hernández, Claudia Rodríguez-Almazán

## Abstract

Laccases belong to the multicopper oxidase family (MCO). Compared with fungal lac-cases, bacterial laccases have high stability and activity under extreme pH and temperature conditions. Because of these characteristics, bacterial laccases have become highly impactful in industrial processes. In the present work, we engineered the *Thermus thermophilus* HB27 laccase (Tth-Lac) by directed evolution. The library had a size of 1.2 x10^5^ colonies, and we screened mutants by malachite green decolorization on solid medium. The results showed that Q36H mutant exhibited higher catalytic activity than Tth-laccase (WT) at 90 °C. Both enzymes showed similar pH profiles, but the activity at 90 °C of Q36H mutant was X-fold higher than that of the wild-type enzyme. In contrast, the kinetic constants were similar for both enzymes. In the voltammetric responses, at 61.5 °C, the Q36H mutant showed increased current, and the T1 copper potential shifted by 100 mV relative to the tth-Lac. These results suggest that the Q36H mutation has high potential for industrial applications because replacing a glutamine (Q) with a histidine (H) at position 36 enhances the enzyme’s structural stability, allowing it to function efficiently under extreme temperature conditions.

## 1. Introduction

Laccases (EC 1.10.3.2) are multicopper oxidases produced by bacteria, fungi, plants, and insects [1–3]. These enzymes were first described by Yoshida in 1883 after isolation from the lacquer tree *Rhus vernicifera* [4]. Because they oxidize a wide range of substrates and remain active under extreme pH and temperature conditions, laccases are attractive for biotechnology, including biobleaching, bioremediation, wastewater treatment, and paper and textile processing [2,5,6]. Although fungal laccases are still the most widely used in industry, bacterial laccases have been receiving special attention because they usually show greater thermal stability and broader tolerance to extreme pH, despite having lower catalytic activities [5,7,8].

Laccases catalyze the oxidation of phenolic compounds and aromatic amines, coupled to the four-electron reduction of oxygen to water [1]. Their catalytic center contains four copper atoms organized into one type 1 (T1), one type 2 (T2), and two type 3 (T3) copper sites. The T1 copper site oxidizes the substrate, and the resulting electrons are transferred through a conserved His–Cys–His pathway to the trinuclear T2/T3 cluster, where oxygen is reduced to water [1,9,10]. This catalytic versatility makes laccases important targets for enzyme engineering.

Over the last two decades, several strategies have been used to improve laccase activity and stability, with directed evolution among the most important [11–13]. This approach allows enzyme optimization without requiring detailed prior structural knowledge. However, only a few bacterial laccases have been extensively engineered, i.e. CotA from *Bacillus subtilis* and CueO from *Escherichia coli*. In CotA, directed evolution has improved substrate specificity, catalytic activity, dye decolorization, solvent tolerance, and acidic stability [14–16]. In CueO, protein engineering has also produced variants with improved redox potential [17].

Among the properties targeted during engineering, redox potential is especially important due to its effect on the oxidizing power and on its effect on the electron transfer to the T1 copper center [18,19]. This parameter influences substrate specificity, catalytic efficiency, and the types of compounds that can be transformed. Although bacterial laccases generally have lower redox potentials than fungal laccases, they remain attractive because of their stability, broad pH tolerance, and ease of production [13,20,21]. In addition, redox mediators can expand their substrate specificity and improve the oxidation of more recalcitrant compounds.

A particularly promising bacterial enzyme is the laccase from *Thermus thermophilus* HB27 (Tth-Lac), first purified and characterized using ABTS and syringaldazine as substrates [22]. Tth-laccase shows high activity near 90 °C and under mildly acidic conditions. It retains about two-thirds of its initial activity after 10 min at 100 °C and has a half-life of 868 min at 80 °C, making it one of the most thermostable multicopper oxidases reported [21,22]. Unlike many laccases, it does not require external copper addition for activity and remains active in a broad interval of temperatures [22,23]. Its T1 copper redox potential of about 0.42–0.53 V falls within the typical for bacterial laccases. However it is strongly affected by temperature and active-site mutations, redox values decreased from 480 mV at 20 °C to 420 mV at 65 °C, showing that temperature substantially influences the redox behavior of this enzyme [24].

The exceptional stability of Tth-Lac has driven interest in industrial and bio electro-chemical applications. It has been used for high-temperature lignin removal in the paper industry and proposed as a catalyst in enzymatic fuel cells because it can reduce oxygen without redox mediators [25,26]. However, its catalytic activity remains relatively low compared with that of other laccases. Therefore, improving its activity and redox potential while preserving thermostability remains an important objective. In this context, the Q36H variant was generated by random mutagenesis to improve catalytic activity and redox potential. Mutants were screened by malachite green decolorization and ABTS oxidation, and the effects of pH, temperature, and electrochemical properties were evaluated. These results support the development of improved thermostable laccases for industrial and bio electrochemical applications.

## 2. Results

### 2.1. Library evaluation and mutant selection

Directed evolution is a protein engineering strategy that generates genetic diversity and identifies protein variants with improved properties through iterative cycles of mutagenesis and screening [27,28]. Directed evolution depends not only on the construction of a mutant library but also on efficient, high-throughput screening methods for rapid identification of variants. To facilitate this technique, enhanced green fluorescent protein (EGFP) is an efficient tool for monitoring library cloning and gene expression, providing a simple and reliable visual marker for identifying recombinant clones. EGFP can be used as a negative selection marker, where disruption or replacement of the EGFP gene results in non-fluorescent colonies that can be readily distinguished from non-recombinant fluorescent colonies, significantly reducing the time and effort required for library screening [29– 31].

The EGFP used in the library construction allowed the identification of colonies with the mutant gene of Tth-laccase (non-fluorescent colonies). To calculate the frequency of mutation, we used the equations (1 and 2) described in the Gene Morph II Random Mutagenesis kit manual.

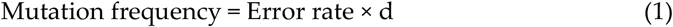

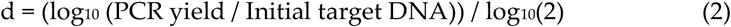

were, mutation frequency are the mutations per kb, error rate is the DNA polymerase error rate (errors • Kb^−1^ • duplication^−1^), and d is the number of duplications during PCR.

these equations show that the mutation frequency increases in proportion to the number of target-molecule duplications during PCR.

An average of 3.4 mutations / Kb was obtained, which corresponds to a low mutation rate. The library had a size of 1.2 × 10^5^ colonies. The diversity of the library was calculated using the Program for Estimating Diversity in Error-prone PCR Libraries (PEDEL) and PEDEL-AA [32]; both programs provided theoretical estimates of the number in the library, considering the gene size, library size, and mutation rate. The sequences analyzed had a 99% probability without insertions or deletions. Each sequence had an average probability of 8% of having a stop codon. For synonymous mutations, it is 2.3 times more probable to detect a non-synonymous mutation than a synonymous mutation. In 20 sequenced colonies, the position of each mutant was identified, and no preferential mutation region was found in the Tth-Lac gene. The region where the mutations were in the 20 colonies was determined. Mutations were found throughout the sequence; none of the areas was favored (Figure 1A).

**Figure 1.**
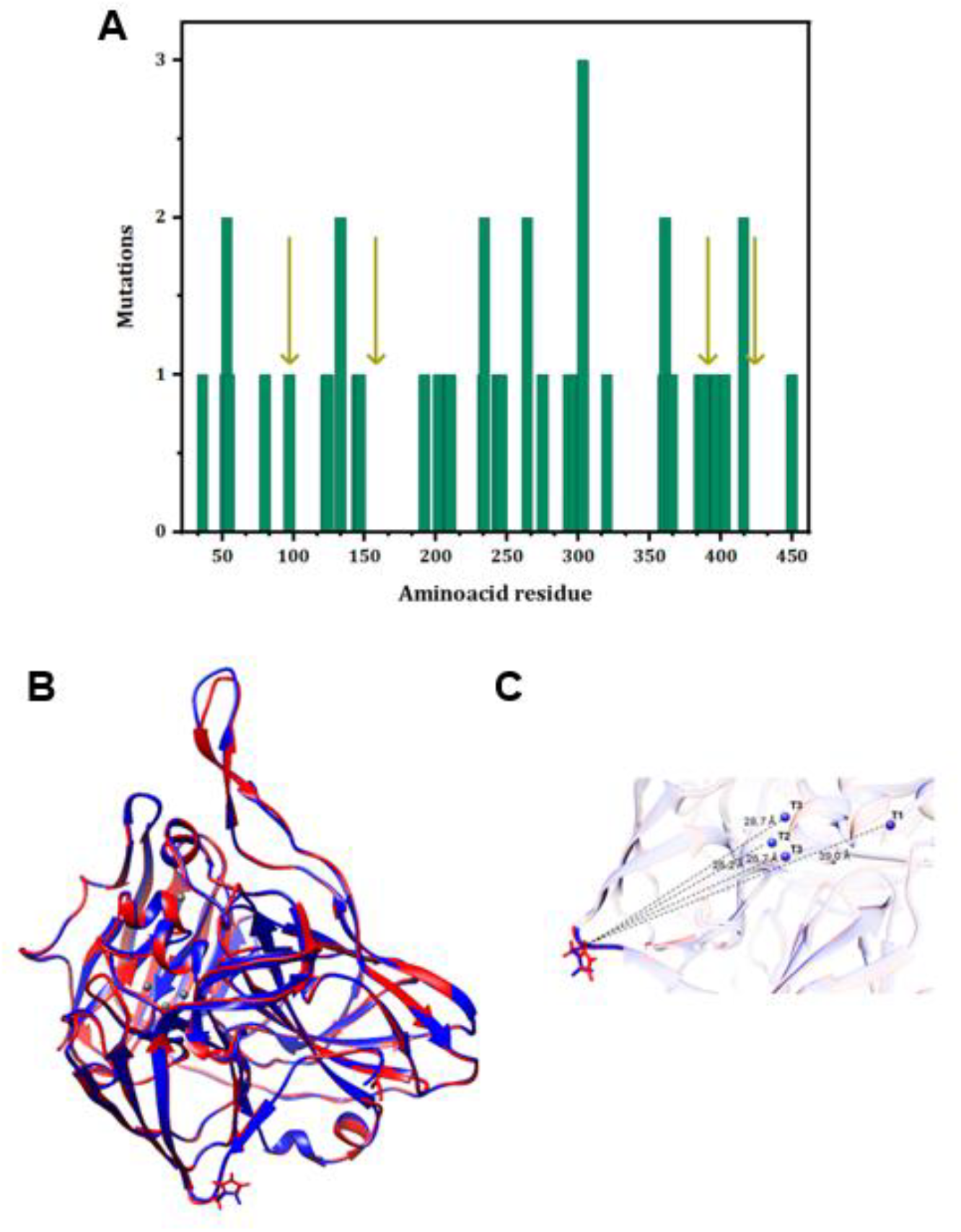
Analysis of the mutant library and structure of Q36H. A. Distribution of mutations along the protein sequence. Each bar represents the average of change in twenty sequenced variants. The arrows indicate the copper coordination position. B. Structural alignment of Tth-laccase (blue) and Q36H mutant (red). These models were from the structure PDB 2xu9 and were used with UCSF Chimera software v.1.17.3 (https://www.cgl.ucsf.edu/chimera/) [33]. C. Distance between the position of H36 and the copper coordination site. The distances between the Q36H mutant and each copper atom are as follows: 39.0 Å to the T1 copper, 26.2 Å to the T2 copper, and 26.7 Å and 28.7 Å to the T3 copper. The distances were calculated with Chimera Software.

In this study, a screening protocol for selecting improved variants from the Tth-Lac mutant library was first standardized. The selection process required overexpression of individual mutants in 96-well plates, as described in the Methods section. Initial screening was performed on solid medium using a dye decolorization assay with seven dyes at pH 5 and 7. As controls, *E. coli* BL21 carrying the empty vector and *E. coli* BL21 expressing pET-22b-Tth-laccase were included. Among the dyes tested, only malachite green at pH 5 showed detectable decolorization (Figure S1). Based on this result, 800 non-fluorescent colonies were randomly selected from the library, overexpressed, and plated on LB agar supplemented with ampicillin and malachite green at pH 5. After incubation at 30 °C for 3 days, 50 colonies, exhibiting greater decolorization than the wild type, were selected.

These variants were subsequently evaluated using an ABTS oxidation assay at 61.5 °C. ABTS oxidation yields a green product detected at 420 nm. The absorptivity increases with higher enzyme activity. Twenty-two variants showing higher oxidation values of ABTS than the wild-type enzyme were selected (data not shown) and further analyzed by measuring their relative activity toward ABTS at 90 °C. Seven variants displaying higher activity were expressed, purified, and characterized by determining their specific activity. Among them, one variant consistently exhibited greater activity than the wild-type enzyme. DNA sequencing revealed a single amino acid substitution, Gln36His (Q36H) (Figure 1 C). To determine the structural location of this mutation, a three-dimensional model was generated using the crystal structure of *T. thermophilus* laccase (PDB ID: 2XU9), and the distance between His36 and the copper coordination site was calculated (Figure 1 D).

### 2.2 Expression, purification and characterization of Tth-laccase and Q36H mutant

The optimal conditions for protein expression and purification were standardized with Tth-Lac. The cultures were grown at different temperatures (18, 25, and 30 °C). tth-Lac and Q36H mutant yields were between 3 and 7 mg per liter of culture, respectively, and at 30 °C, 5 mg per liter of culture medium was obtained. However, the protein with the highest activity was the one expressed at 30 °C (Figure S3); this condition was used to express Tth-Lac and the Q36H mutant. Tth-Lac and the Q36H mutant were expressed as a holoenzyme (with copper in the culture medium). The purification protocol consisted of a chromatography step on a CM-Sepharose ion-exchange column. The protein eluted from the buffer fraction at 40-70 mM NaCl (Figure S2). The purified proteins were analyzed under denaturing conditions by electrophoresis (12% acrylamide SDS-PAGE gel), in which a band was observed at the expected molecular weight (Figure 2A).

**Figure 2.**
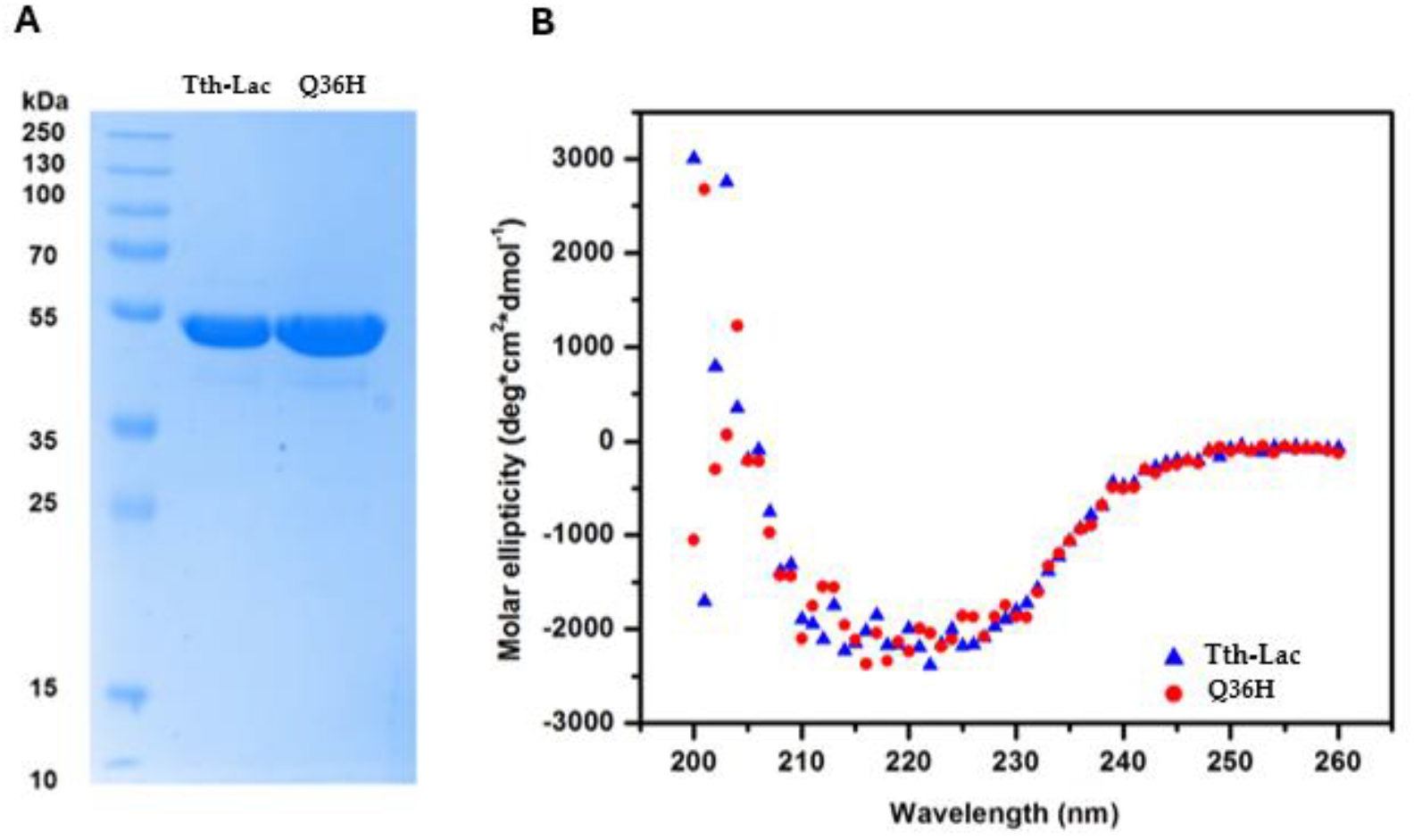
Characterization of Tth-Lac and Q36H mutant. Tth-Lac and Q36H mutant purified by CM-Sepharose column. A. SDS-PAGE electrophoresis in 12% acrylamide gel. B. CD spectra of Tth-Lac (blue) and Q36H mutant (red) at 25 °C, 200 μg / mL protein concentration, in 20 mM phosphate buffer, pH 4.5.

The far-UV CD spectra of Tth-Lac and Q36H mutant were analyzed to identify a possible change in secondary structure. Circular dichroism signals of Tth-Lac and Q36H mutant showed the same negative band at 217 nm. This result indicated no change in secondary structure due to the mutation (Figure 2B).

The kinetic parameters were measured at 61.5 °C to compare with previously reported values for the same enzyme [34]. The Q36H mutant and the Tth-Lac differ in the kinetic parameters determined at 61.5 °C using ABTS as the substrate (Table 1). Compared with Liu *et al*. (2011), our results show that Tth-Lac and Q36H exhibit considerably higher values in catalytic parameters than the previously reported Tth-Lac. However, it is important to consider the differences in experimental conditions. Liu et al. used 10 µg of enzyme and 100 mM copper sulfate in the activity assay, whereas here we used 1 µg of enzyme and 1 mM copper sulfate. These differences can influence enzymatic activity and make a direct comparison between the two studies difficult. Therefore, the results suggest that the Tth-Lac and Q36H enzymes may exhibit even higher catalytic activity than the enzyme reported by Liu *et al*. (2011).

**Table 1.**
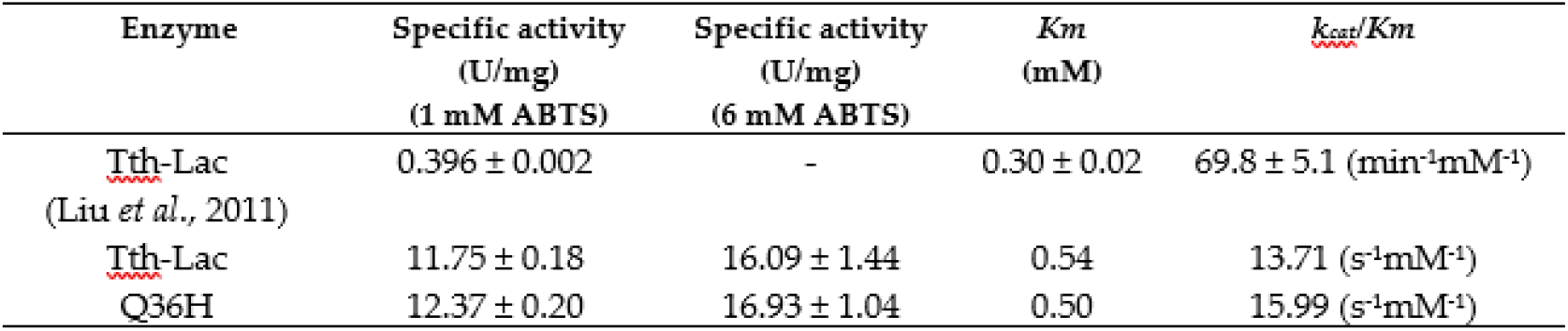
Kinetic parameters of Tth-Laccase and Q36H mutant at 61.5 °C, pH 4.5 and ABTS as substrate.

| Enzyme | Specific activity<br>(U/mg)<br>(1 mM ABTS) | Specific activity<br>(U/mg)<br>(6 mM ABTS) | $K_m$<br>(mM) | $k_{cat}/K_m$ |
| --- | --- | --- | --- | --- |
| Tth-Lac<br>(Liu *et al.*, 2011) | $0.396 \pm 0.002$ | - | $0.30 \pm 0.02$ | $69.8 \pm 5.1$ (min<sup>-1</sup>mM<sup>-1</sup>) |
| Tth-Lac | $11.75 \pm 0.18$ | $16.09 \pm 1.44$ | 0.54 | $13.71$ (s<sup>-1</sup>mM<sup>-1</sup>) |
| Q36H | $12.37 \pm 0.20$ | $16.93 \pm 1.04$ | 0.50 | $15.99$ (s<sup>-1</sup>mM<sup>-1</sup>) |

The specific activity of both enzymes was measured from 61.5–90 °C in order to determine whether the change of Gln to His at position 36 affected activity at temperatures above 61.5 °C. In this experiment, the mutant is approximately 20 U/mg more active than the Tth-Lac with 1 mM ABTS. In contrast, at extreme temperature (90°C), the difference is approximately 5 U/mg under saturating conditions; the mutant still performs better than the wild type, but the improvement is smaller.

Tth-Lac and the Q36H mutant showed difference at 90 °C (Figure 3A), indicating that the Q36H substitution favored catalytic activity at extreme temperatures. Because lac-cases can catalyze at extreme pH values, the specific activity was measured over the pH range of 3–6. These experiments were performed in the absence of copper in the reaction medium. Based on observations in our laboratory and previous reports, copper precipitates at pH ≥ 6, which interferes with activity measurements. The results obtained for enzymatic activity at different pH values show that both enzymes exhibit the same behavior. A maximum activity was observed at pH 4.5 (Figure 3B).

**Figure 3.**
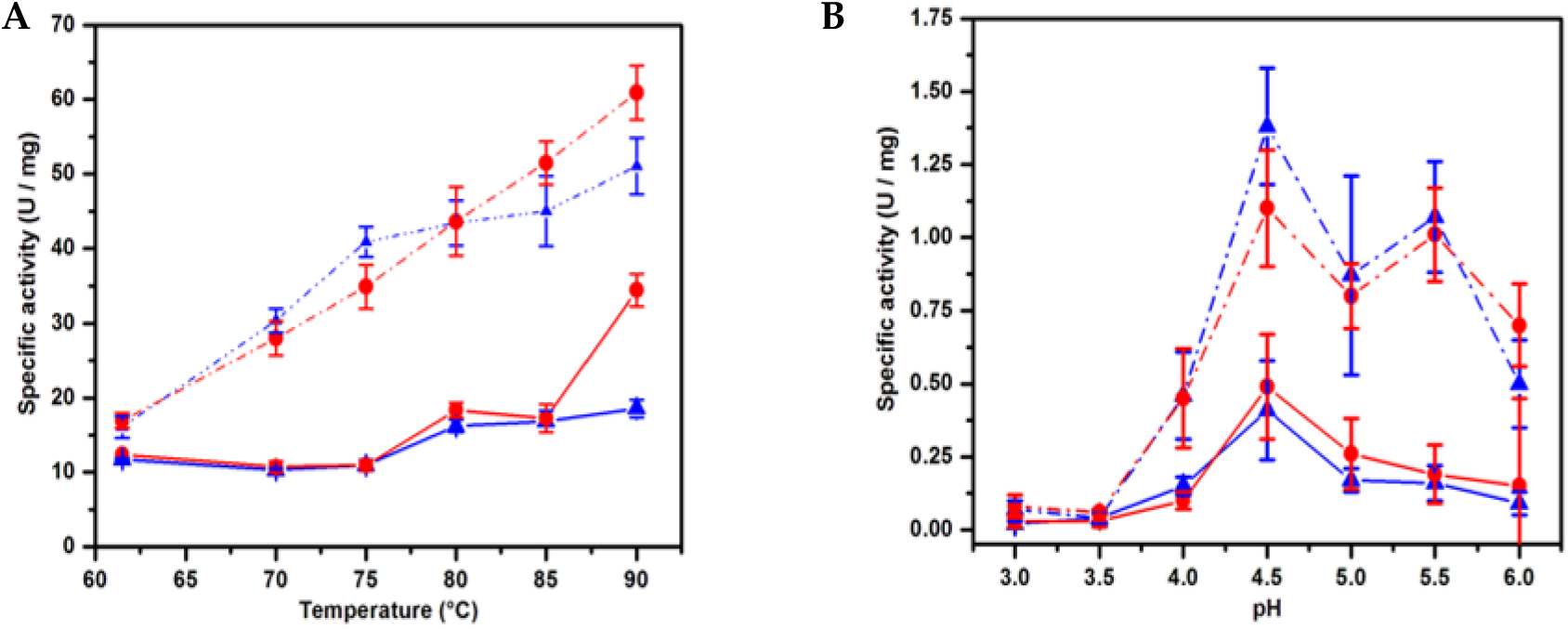
Effect of temperature and pH on the activity. Catalytic activity at different temperatures (A) and pH (B). The experiments were conducted with 1μg of enzyme, 61.5 °C, 1 mM ABTS as substrate, 20 mM sodium acetate buffer, in the case of the activity at different temperatures, the reaction was run at pH 4.5. The solid line represents catalytic activity using 1 mM ABTS; the dashed line represents the catalytic activity with 6 mM ABTS. Tth-Lac is blue color, and Q36H mutant is red color.

**Figure 4.**
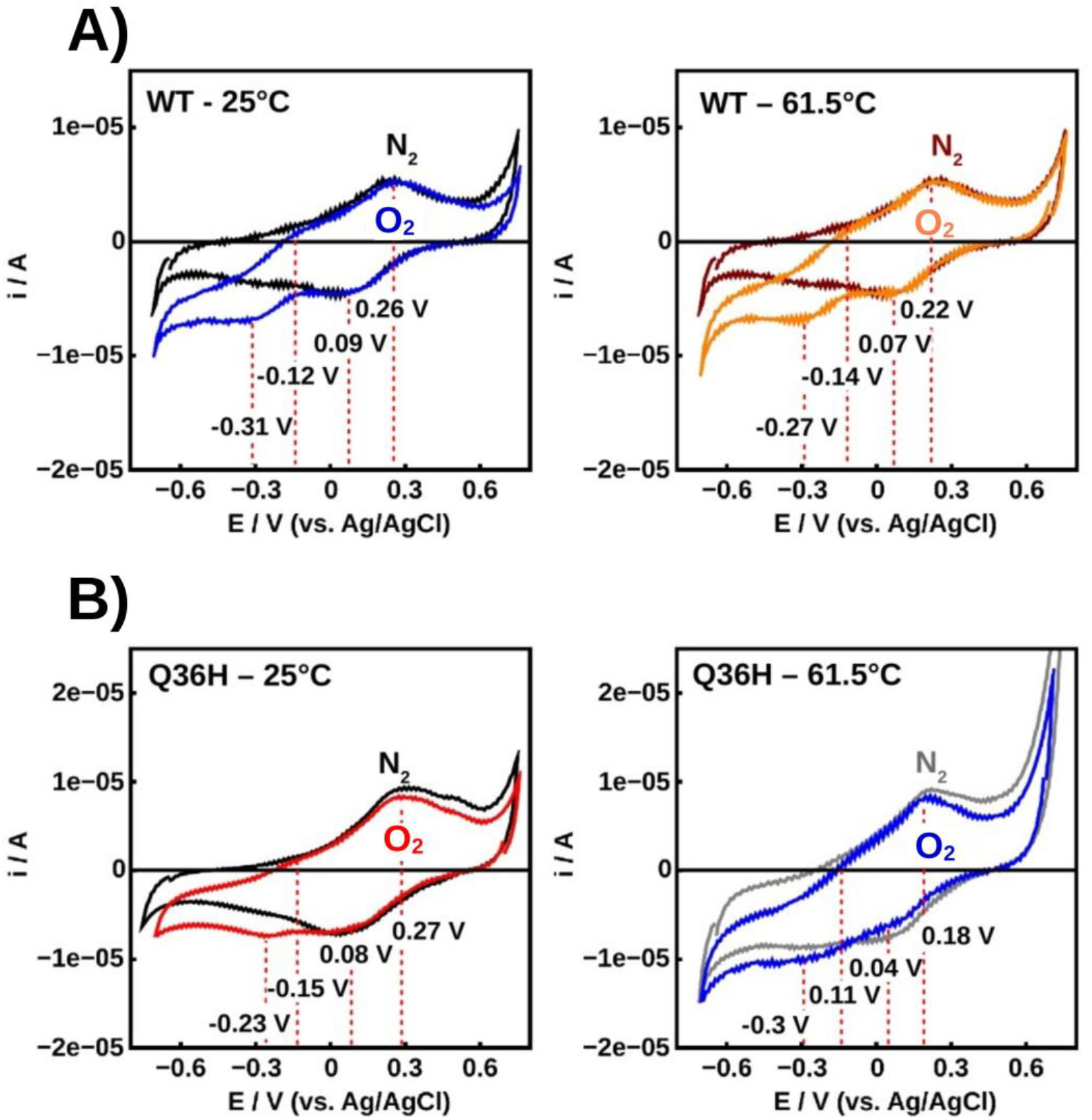
Voltametric response of A) Tth-Lac (WT) and B) Q36H mutant. The responses were measured at 25 °C and 61.5 °C, in 0.1 M sodium acetate buffer, 0.1 M KNO_3_ pH 5, at 0.005 V/s, in anaerobic (N_2_) and aerobic (O_2_) atmospheres.

The voltammetric responses of the Q36H enzyme at 25 °C showed a change upon switching the atmosphere from nitrogen to air. Unlike the wild type enzyme, a reduction process can be observed, with an inflection at a potential of −0.23 V. In the case of copper T1, the potential response is lower than that of the copper T1 from the wild enzyme. When the experiments were conducted at 61.5 °C, there was a noticeable increase in current, unlike the wild enzyme. The copper T1 response shows a 100 mV displacement in the negative direction.

The analysis of voltammetry of the wild enzyme and Q36H mutant showed that at 25 °C, the enzymatic activity can be clearly observed when the atmosphere passed from nitrogen to air, with an increase in the reduction current of 0.0 V and −0.23 V, respectively, and with an inflection in a potential value of −0.31 V only for the case of the wild enzyme. When the experiment was measured at 61.5 °C, Tth-laccase did not show an appreciable change in current or peak potential values. Unlike the wild enzyme, a reduction process is observed, with an inflection at −0.23 V. In the case of T1, the peak potential is at a lower potential compared to the respective TG1 of wild type. When the temperature rose to 61.5 °C, there was a noticeable increase in current, unlike the wild enzyme. It was further observed that the peak potential of T1 was displaced by 100 mV in the negative direction. This data could explain the increase in the enzymatic activity of the Q36H mutant at this temperature.

## 3. Discussion

Directed evolution is one of the most successful methods for improving the function and stability of enzymes without structural knowledge. However, identifying improved variants depends on the diversity generated, the quality of the mutant library, and the sensitivity of the screening method. The Tth-Lac mutant library showed a mutation frequency within the optimal range for directed evolution, and this study reported, for the first time, a solid medium with a dye screening strategy for engineering laccase.

The identification and characterization of the Q36H mutant located in an unexplored region provides new insights into the structural determinants for Tth-Lac activity. This finding is particularly relevant because enhancing activity while maintaining thermostability remains one of the principal challenges in the protein engineering of extremophilic enzymes for industrial applications, and it highlights previously unexplored regions that may contribute to catalysis.

Circular dichroism (CD) analysis showed that the Q36H substitution did not alter the overall secondary structure of Tth-Lac, as demonstrated by the nearly identical spectra of the wild-type and mutant enzymes. The spectra of wild-type Tth-Lac and the Q36H variant were nearly identical.

Likewise, kinetic parameters determined at 61.5 °C, did not differ significantly. The improved catalytic activity of Q36H at 90 °C is likely due to subtle local changes in the structural or electronic environment of the enzyme rather than global conformational re-arrangements. The catalytic activity of Q36H at high temperature is consistent with previous studies showing that mutations identified by directed evolution are often located outside the active site and improve enzyme performance by modulating protein dynamics, long range interactions, or electron transfer pathways while preserving the catalytic mechanism [35,36]. In this context, histidine possesses an imidazole side chain capable of establishing hydrogen bonds; the imidazole side chain provides the functional versatility needed for such dynamic tuning, as it can form hydrogen bonds, coordinate metal ions, and participate in proton-transfer networks [37,38]. Consequently, changing amino glutamine in position 36 with histidine may improve catalytic conformations at high temperature without perturbing the native fold.

Elevated temperatures destabilize proteins via denaturation and accelerate nonenzymatic chemical alterations, notably the deamidation of asparagine and glutamine [39– 41]. This modification introduces a negative charge that disrupts protein interactions and function, with implications for biological aging, disease, and pharmaceutical shelf-life. Because deamidation rates depend on local structural context and neighboring residues, susceptibility varies widely across the protein. While this process directly drives temperature induced loss of function, past mutational research has primarily focused on mapping these consequences rather than optimizing stability.

Although Tth-Lac contains numerous glutamine (Gln) and asparagine (Asn) residues throughout its structure, non-enzymatic deamidation depends strictly on local chemical environments and specific spatial orientations [42,43]. For instance, deamidation leading to inactivation relies on neighboring residues, such as the Thr-Asn-Glu sequence in cytochrome c [44,45]. In this sense, the Q36H mutant possesses the requisite orientation and microenvironment to undergo deamidation, which likely contributes to its altered catalytic behavior.

The low catalytic efficiency of Tth-Lac has hindered its industrial use despite its intrinsic thermostability. Thus, the Q36H mutant has greater activity than Tth-Lac and is 400-fold higher than previously reported values [46], highlighting the combined benefits of enzyme engineering and optimized expression, purification, and copper supplementation, as described in other studies [23].

In the case of voltammetric results indicated that the Q36H mutation modified the electrochemical behavior of the enzyme at 61.5 °C; the mutant exhibited an increase in reduction current and a shift of nearly 100 mV toward more negative potential at the Cu T1 center, compared to Tth-Lac. This finding is significant because, in laccases and other multicopper oxidases, the T1 site serves as the primary entry point for electrons from the substrate and is functionally coupled to the T2/T3 trinuclear cluster, where oxygen is reduced to water. Relatively small variations in Cu T1 potential can alter substrate oxidation and electron transfer to the trinuclear cluster [14,16]. In CotA laccase, mutations in the T1 environment shifted the redox potential by magnitudes of 60 to 100 mV, as well as spectroscopic, catalytic, and geometric changes at the metal center. In another study, substituting the axial ligand in CotA increased the potential by approximately 100 mV while preserving the general geometry of the T1 site and protein folding, demonstrating that changes in the copper microenvironment are sufficient to reprogram its redox properties [47]. Other studies have demonstrated that rational engineering can simultaneously improve redox potential, catalytic activity, and stability, supporting the idea that well-positioned mutations can optimize multiple functional traits at once [21,48,48,49]

These results contribute to understanding the structural determinants influencing bacterial laccase activity for future studies of directed evolution to optimize enzymes with applications in industrial processes.

## 4. Materials and Methods

### 4.1 Expression and purification

Gene cloning of Tth-Lac was performed as described previously [50], electrocompetent *Escherichia coli* BL21 Star™ (DE3) cells were transformed and grown in Luria-Bertani (LB) medium. For expression, 50 mL of LB broth containing 100 μg/mL ampicillin was inoculated with a colony and incubated for 14 h at 37 °C and 250 rpm. After 1 L of LB medium was inoculated, the initial optical density (O.D.) at 600 nm was 0.05. The culture was incubated at 37 °C and shaken at 250 rpm. Protein expression was performed at an O.D.600 nm equal to 0.6, with isopropyl-β-D-thiogalactoside (IPTG) (Thermo Scientific) and CuSO4 (Sigma Aldrich), both at a final concentration of 1 mM, at 30 °C, and 250 rpm. After 4 hours, the culture was maintained at the same temperature for an additional 12 hours without stirring. Cells were harvested by centrifugation at 3,000 × g for 30 min at 4 °C. Cell lysis was performed in 50 mL of 50 mM 3-(N-Morpholino) paropanelsulfonic acid (MOPS) buffer (Sigma-Aldrich), pH 7.4, with 1 mM phenylmethylsulfonyl fluoride (PMSF) (Sigma Aldrich). For cell lysis, an ultrasonic sonicator (Cole-Parmer) was used for six cycles (35% amplitude, 59 pulses, 40 s). The sample was centrifuged at 35,000 × g for 40 min at 4 °C. To remove all labile proteins from *E. coli*, the supernatant was incubated at 65 °C for 30 min in a Lab-Line thermobath (Thermo Scientific), then the sample was centrifuged at 35,000 × g for 20 min at 4 °C. Protein purification was done in 10 mL of an ion-exchange cationic resin (CM-Sepharose, GE Healthcare), equilibrated with 50 mM MOPS, pH 7.4. The sample was loaded onto the column at 0.3 mL/min, and the chromatography was performed at 1 mL/min. A NaCl gradient was used to elute the protein (0 mM, 20 mM, 40 mM, 70 mM, 100 mM, and 500 mM). The fractions collected were analyzed by denaturing SDS-PAGE (12%) and stained with Coomassie blue (Sigma-Aldrich). The fractions containing laccase were concentrated by ultrafiltration (Amicon Ultra-3, Millipore) using a 30-kDa cut-off membrane. Then, the protein was dialyzed in 20 mM sodium acetate buffer, 0.1 mM CuSO4, pH 6.0 for 48 h, at 4 °C; subsequently dialyzed against 20 mM sodium acetate buffer, 0.1 mM CuSO4, pH 4.5 for 24 h at 4 °C. To eliminate aggregates, the sample was centrifuged at 8,000 × g for 10 min at 4°C. Protein concentration was determined by absorbance at 280 nm. The molar extinction coefficient of Tth-laccase is 46,410 M-1cm-1. Finally, the protein was stored at 4°C until use.

### 4.2 Circular Dichroism

This analysis was performed using a JASCO J-715 Spectropolarimeter (JJASCO Inc.) with a filled cell holder in a 0.1 cm cell. Each filtered sample contained 500 μg protein/mL in 20 mM sodium phosphate, pH 6.0. Spectral scan was performed from 198 to 260 nm, with intervals of 20 nm/min at 25 °C.

### 4.3 Laccase activity and kinetics parameters

To measure the enzymatic activity at temperatures below and equal to 61.5 °C, the Cary 60 UV-Vis (Agilent Technologies) spectrometer was used, with a PCB 1500 Water Peltier System (Agilent) temperature controller, while for temperatures above 62 °C, the DU 650 spectrometer (Beckman) adapted to the High-Performance Temperature Controller (Beckman) was used. Enzymatic activity was measured according to the method previously reported (Liu 2011). The reaction mixture consisted of 20 mM sodium acetate buffer, pH 4.5, 1 mM CuSO4, 2,2′azino-bis (3-ethylbenzothiazoline-6-sulfonic acid) (ABTS) (ε420 = 36,000 M-1 cm-1) as substrate, and 1 μg/mL (20.35 pM) of protein. Specific activity was measured with 1 mM ABTS at 61.5 °C. To calculate *Km* and *kcat*, different concentrations of ABTS (0.75 – 6 mM) were used at 61.5 °C. The activity was determined at different temperatures and pH with 1 μg/mL of protein, 1 mM ABTS, and, in the case of pH, at 61.5 °C. In all experiments, it was expected that the buffer in the cell would take three minutes to reach the same temperature as the cell holder. As a control, the reaction medium was used without the enzyme. All catalytic activity measurements were performed in quadruplicate.

### 4.4 Library creation

The library was built using the random mutagenesis technique with Gene Morph II Random Mutagenesis kit using the Tth-laccase gene as templated. To increase the percentage of mutations, 10 ng of PCR product obtained was subjected to a second round of PCR. To remove remaining of DNA polymerase attached, the PCR product was digested with proteinase K (Roche) 1 U / 100 ng of DNA, 1 h at 37 °C [51]. The vector digested with NdeI and EcoRI restriction enzymes and purified was incubated with shrimp alkaline phosphatase (New England). The reaction was in a molar ratio of 1:3 (vector/insert) with 1 U of T4 DNA ligase (New Enland), 0.05 mM ATP, 16 n at 18 °C. The library was cloned into pET22b (+) and transformed into electrocompetent *E. coli* BL21 (DE3) strain. To select the constructions with mutantants, the green fluorescent protein (EGFP) was used as a reporter molecule. When EGFP is expressed in *E. coli*, it shows fluorescence upon exposure to UV light, this protein has been used in ligation reactions, since it is possible to select the correctly linked constructs based on those that do not exhibit fluorescence. EGFP gene was in the vector pJOQ, so, it was amplified by PCR using primers designed with the restriction sites NdeI and EcoRI (EGFP-NdeI: 5’ AGGAGATATA**CATATG**GTGAGCAAG 3’, EGFP-EcoRI: 5’ACGGAGCTC**GAATTC**CTTGTACAGC 3’), was cloned into pET22b (+) and transformed into *E. coli* BL21 (DE3). The cells were plated in LB medium with 200 μg/mL ampicillin, and incubated at 30 °C, for 16 h. To determine the rate of mutagenesis, twenty-one colonies were selected and their DNA sequenced.

### 4.5 Selection of mutants

For the selection assay, the protein had to be overexpressed. Selected colonies (non-fluorescent) were transferred to 96-well plates with 200 μL of LB/ampicillin medium and incubated for 12 h at 37 °C at 250 rpm. Then, 100 μL of culture were placed on a separate plate, and 100 μL of glycerol (80 stock) were added to each well, which were then stored at −80 °C. Then, 10 μL of the same culture was used to inoculate 190 μL of LB/ampicillin medium. These cultures were grown for 2.5 h at 37 °C and 250 rpm, after which protein expression was induced with 1 mM IPTG and 1 mM CuSO4. The cultures were incubated for 4 h at 30 °C and 250 rpm, then incubated at 30 °C without stirring for 12 h. The cells were recovered by centrifugation at 2000 × g for 40 min at 4 °C. The cells were analyzed using two selection methods. As the first method, decolorization of different colorants was evaluated. The selected and expressed colonies were seeded in solid LB / ampicillin culture medium at pH 5 and 7, adjusted with 1 N HCl and 1 N NaOH. The decolorization assay was performed on different dyes, each prepared as a solution containing 0.1 g of dye in 1 mL of culture medium, using a 1:10,000 dilution of each dye from Sigma Aldrich. Colonies were seeded in dye containing media and incubated at 30 °C for 72 h. As a second method of selection, the relative activity was analyzed for 800 variants using ABTS as the substrate. Cells were resuspended in 20 μL of 50 mM MOPS, pH 7.4, containing 1 mM PMSF. To lyse the cells, 1 μL of B-Per detergent (Roche) was used; the mixture was incubated at 30 °C for 1 h. To remove cell debris, the plates were centrifuged at 2000 × g for 40 min at 4 °C; the supernatant was transferred to a new 96-well plate to evaluate the enzymatic activity. Lac-case overexpression was analyzed by denaturing SDS-PAGE gel electrophoresis (12 % acrylamide) from 20 colonies. The ABTS oxidation reaction was carried out at 60 °C for 20 min in a Thermomixer (Eppendorf). For the reaction, 180 μL of 20 mM sodium acetate buffer (pH 4.5), 1 mM CuSO4, and 3 mM ABTS were added. The reaction product was detected at 420 nm, and colonies with higher optical density than the controls (wild-type enzyme) were selected. As a third method of selection, the variants with higher activities in the two methods mentioned above were purified as wild-type enzymes, and their specific activity was measured at 90 °C using ABTS as the substrate, under conditions identical to those described above for the wild-type enzyme. These variants were expressed in a 96-well plate in 200 μL of LB /ampicillin medium under the same conditions as the library. Cell lysis was performed with B-Per (Roche); the extracts were incubated at 60 °C for 20 min and centrifuged. Finally, the mutant and wild-type enzymes were expressed in *E. coli*.

### 4.6 Enzymatic activity at different pH and temperatures

In order to determine the activity at different pHs, the enzymatic reactions were measured at pH 4-5.5, at 61.5 °C, 1 μg (20.35 pM) of protein, 1 mM ABTS, 20 mM sodium acetate buffer, while at pH 6, the reaction media contained 20 mM 1,4-piperazi-nediethanesulfonic acid (PIPES) (Sigma Aldrich) buffer. The reaction was initiated by adding 6 mM ABTS as the substrate. Activity was followed by the increase in absorbance at 420 nm as a function of time in an Agilent Cary 60 spectrophotometer. The effect of temperature on the activity of the enzymes was measured using 20 mM sodium acetate buffer (pH 4.5), 1 mM CuSO4, 1 μg (20.35 pM) of enzyme, and 1 mM ABTS as the substrate. The variable temperatures were 61.5 °C, 70 °C, 80 °C, 85 °C, 90 °C and 92 °C.

### 4.7 Cyclic Voltammetry experiments

We conducted electrochemical studies using a carbon film electrode (CFE) prepared from an isopropanol solution containing 16 mg/mL carbon material and 1% Nafion®. These components were mixed via sonication for 30 minutes to form the film. To enhance the conductivity of the CFEs and enable covalent enzyme coupling, gold was deposited using a solution of 1.3 mM HAuCl_4_ / 0.4 M K_2_HPO_4_/0.2 M KCl (pH 7) by applying a potential pulse of −0.15 V for 60 seconds. Next, add 3 μL of 0.2 M 4-aminothiophenol solution in absolute ethanol. After 20 minutes, the electrode was immersed in 0.1 M HClO_4_, and—to promote 4-aminothiophenol dimerization—seven cyclic voltammetry cycles were performed between 0.1 and 0.8 V (vs. Ag/AgCl) at a scan rate of 0.02 V/s in the oxidative direction. Finally, the electrode was rinsed and dried at room temperature. Covalent attachment of the enzyme to the electrode was promoted using a mixture of 4 mg/mL N-(3-dimethylaminopropyl)-N’-ethylcarbodiimide (EDC) hydrochloride / N-hydroxysuccinimide (NHS) and 400 μg/mL of the enzyme in 0.1 M H_2_PO_4_^−^/H_2_PO_4_^−2^ buffer (pH 5); the mixture was incubated for 1.5 h at 4 °C to control the autohydrolysis of the EDC and NHS precursors. Afterward, the electrodes were washed with buffer. To promote the orientation of the catalytic site (T2/T3 copper) toward the electrode, covalent bond formation was induced using 4-mercapto-N-phenylquinone diimine (MPD), a mixture of N-ethyl-N’-(3-dimethyl-aminopropyl)carbodiimide (EDC) hydrochloride and N-hydroxysuccinimide (NHS) at 4 mg/mL, and 400 μg/mL of enzyme in 0.1 M H_2_PO_4_^−^/HPO_4_^2-^ buffer (pH 5) for 1.5 h at 4 °C. Subsequently, the electrodes were washed with buffer.

### 4.8 Statistical analysis

All experiments were carried out at least by triplicate. The data were analyzed with the Origin Pro software program (version 9.0).

## Supplementary Materials

The following supporting information can be downloaded at https://www.mdpi.com/article/doi/s1, **Figure S1**. Selection of mutants via malachite green decolorization. **Figure S2**. Specific activity of Tth-Lac under different overexpression conditions. **Figure S3**. Purification of Q36H mutant.

## Author Contributions

C.R.-A., B.M.-Z., P.G.-C., M.M.-H., and G.H.-M. performed the conceptualization, designed the experiments, and performed the experiments. C.R.-A., B.M.-Z., P.G.-C., M.M.-H., G.H.-M. and L.R.-S. analyzed the data. C.R.-A., B.M.-Z., P.G.-C., M.M.-H., and G.H.-M. wrote the paper. All authors have read and agreed to the published version of the manuscript.

## Data Availability Statement

The authors acknowledge all the support provided by the Institute of Biotechnology and the Institute of Cellular Physiology, both Universidad Nacional Autónoma de Mexico (UNAM), for the facilities granted in various laboratories to carry out experiments. The authors acknowledge the technical assistance of María Concepción José Núñez, Juan Carlos Campero Basaldua, and Juan Manuel Barbosa Castillo, special acknowledgment to Alicia Gonzalez Manjarrez all from IFC, UNAM., and Sequencing and synthesis Unit of DNA IBT-UNAM.

## Abbreviations

The following abbreviations are used in this manuscript:

ABTS: 2,2’-azino-bis(3-ethylbenzthiazoline-6-sulfonic acid)
CueO: laccase from *Escherichia coli*
CotA: laccase from *Bacillus subtilis*
EGFP: enhanced green fluorescent protein

## Disclaimer/Publisher’s Note

The statements, opinions and data contained in all publications are solely those of the individual author(s) and contributor(s) and not of MDPI and/or the editor(s). MDPI and/or the editor(s) disclaim responsibility for any injury to people or property resulting from any ideas, methods, instructions or products referred to in the content.

