## Supplemental Figures S1, S2,S3 for "Directed evolution of laccase from *Thermus thermophilus* HB27 reveals a variant with enhanced high-temperature catalytic activity and electrochemical properties"

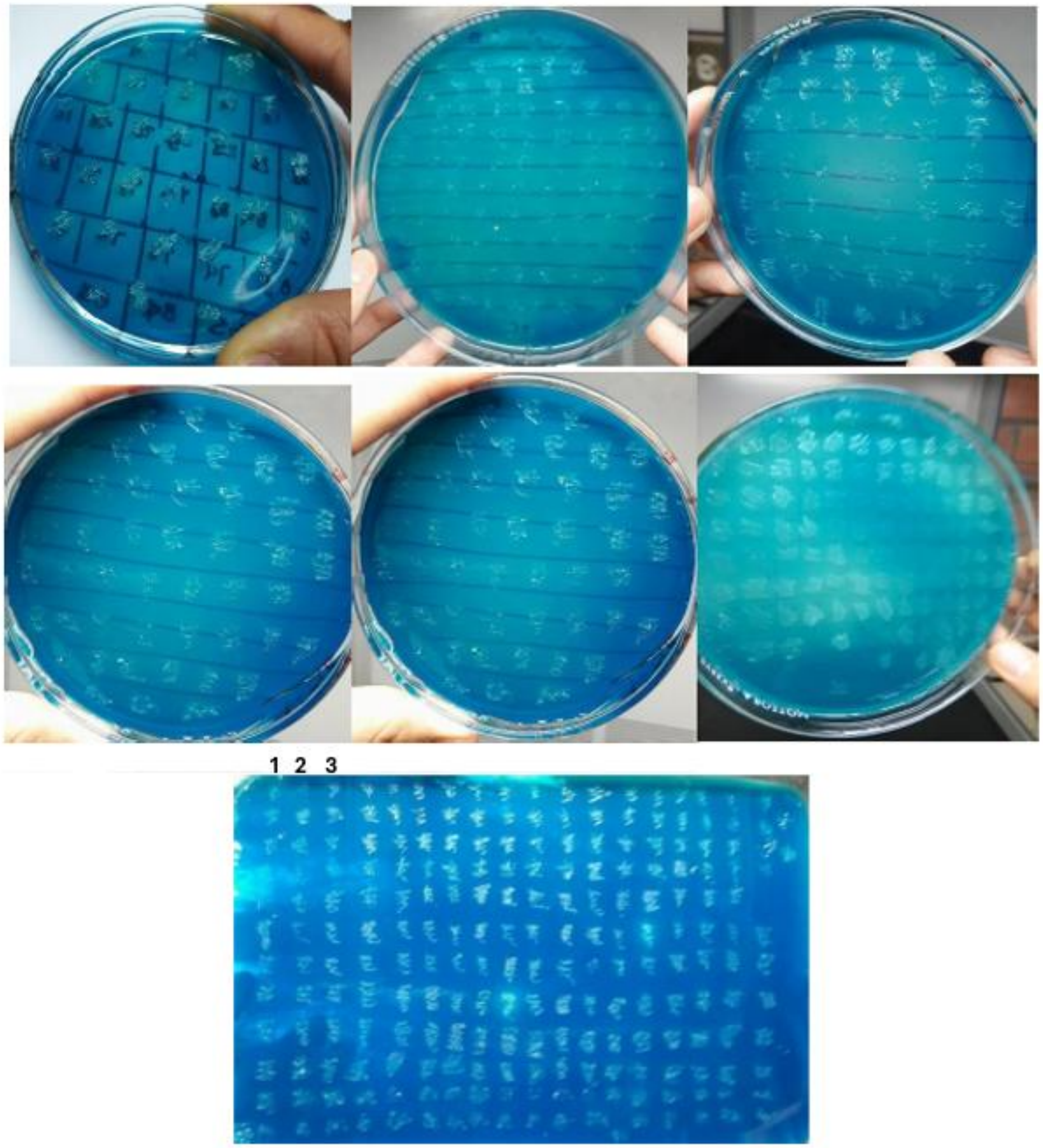


**Figure S1**. Selection of mutants via malachite green decolorization. Some colonies analyzed in the selection are shown stained with dye. **1.** *E. coli* BL21, **2**. *E. coli* BL21 + pET22b, **3.** *E. coli* BL21 + pET22b + Tth-Lac. LB culture medium, 200 μg/mL ampicillin, pH 5.0, and malachite green.

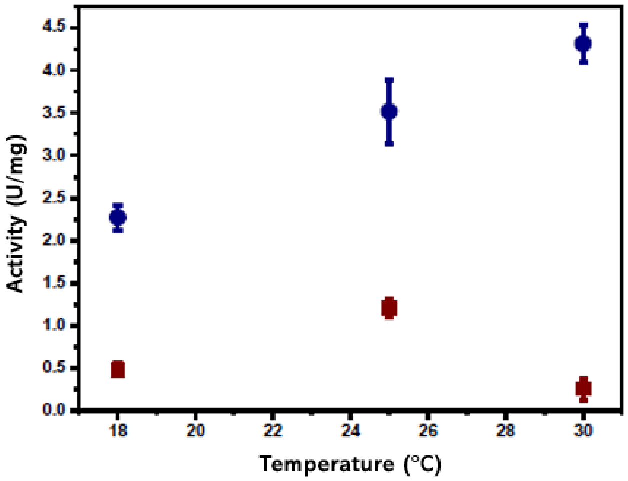


**Figure S2.** Specific activity of Tth-Lac under different overexpression conditions. In the figure, the red square represents cultures without agitation following overexpression, and the blue circle represents cultures subjected to 4 hours of agitation after overexpression.

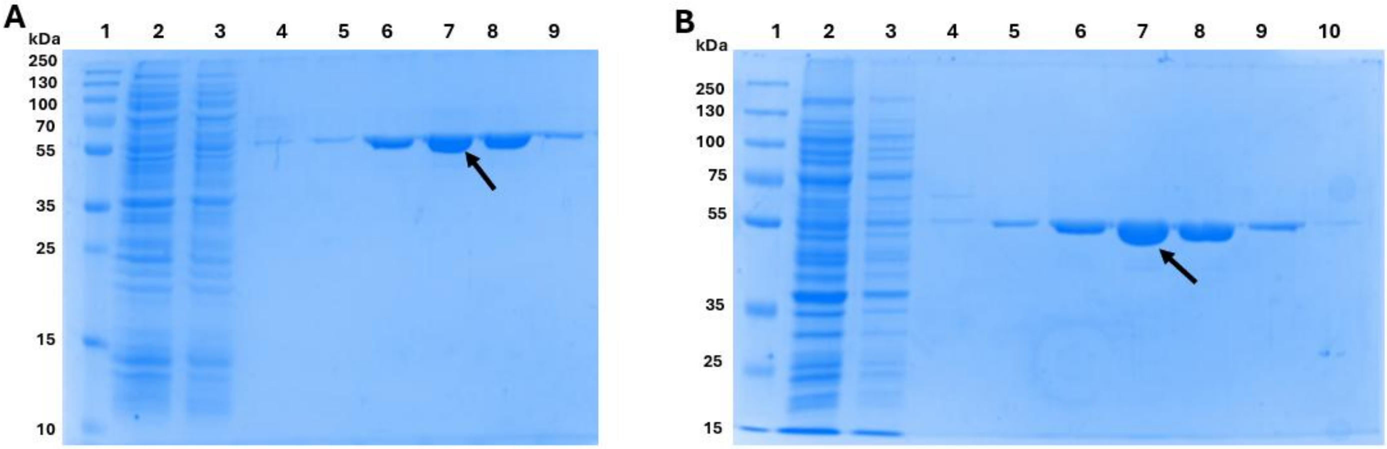


**Figure S3**. Purification of Tth-Lac and Q36H mutant. Line 1, ladder. Line 2, crude extract. A) Tth-Lac and B) Q36H mutant. Protein eluted with NaCl gradient, Line 3, 20 mM NaCl. Line 4 to 5, 40 mM NaCl; lines 6 to 9, 70 mM NaCl; and line 10, 100 mM NaCl. SDS-PAGE, 12% acrylamide, and Coomassie stain. The arrow points to the mutant.
